# ATR facilitates the degradation of Api5 through the ubiquitin-proteasome pathway via FBXW2 to regulate apoptosis upon DNA damage

**DOI:** 10.1101/2021.08.08.455545

**Authors:** Virender Kumar Sharma, Sehbanul Islam, Janhavi Borkar, Sudiksha Mishra, Debiprasad Panda, Manas K Santra, Mayurika Lahiri

## Abstract

Apoptosis inhibitor 5 (Api5) is an inhibitor of apoptosis, which is found to be upregulated in several cancers and promotes invasion as well as metastasis. Over-expression of Api5 is positively co-related with poor survival of cancers and inhibition of DNA damage induced apoptosis in cancerous cells. Acetylation at lysine 251 (K251) on Api5 facilitates the stability of the protein and thus functionally provides resistance to cancer cells against chemotherapeutic or anti-cancerous agents. However, the regulation of Api5 upon DNA damage is not yet known. In this study, we demonstrate that Api5 undergoes degradation following DNA damage via the ubiquitin-proteasome system. Upon DNA damage, ATR was observed to phosphorylate Api5 at serine 138 which led to the cytoplasmic localisation of Api5. The E3-ubiquitin ligase, SCF-FBXW2 ubiquitinates Api5 leading to its proteasomal degradation.

## Introduction

Apoptosis inhibitor 5 (Api5) is a ubiquitously present protein which was initially discovered as a negative regulator of apoptosis under stressed conditions (1). Api5 protein is comprised of α-helical repeats and has three main domains: LxxLL (leucine repeat domain) is responsible for maintaining the stability of the protein, a putative LZD (leucine zipper domain) that is generally known to bind to chromatin and NLS (Nuclear Localisation Sequence) that is responsible for the transport of the protein to the cytoplasm from the nucleus (2). Depending upon the cellular context, Api5 has been shown to engage in its anti-apoptotic function through interaction and regulation of multiple proteins. As mentioned earlier, Api5 was found to inhibit apoptosis upon serum starvation (1), however, later it was observed that Api5 could also play a role in inhibiting apoptosis during various stresses like viral infections, endotoxins and DNA damage. Api5 regulates Acinus-mediated DNA fragmentation due to acute DNA damage-induced apoptosis. Api5 binds with Acinus and prevents its activation (3). In human osteosarcoma cells, ectopic expression of Api5 was found to reduce E2F1-mediated apoptosis (4). Api5 has also been observed to aid in the degradation of Bim, a pro-apoptotic protein (5). It has also been reported that Api5 inhibits the transcription of APAF, one of the main components of the apoptosome complex. A recent study reported the interaction between Api5 and the zymogen, caspase 2. Upon binding with caspase 2, Api5 prevents its activation (6).

Acetylation at lysine 251 (K251) is the only reported post-translational modification of Api5 (2). This K251 acetylation provides stability to Api5. It has been observed that acetylation at lysine 251 on Api5 by p300 histone acetyl transferase stabilises while HDAC1-mediated de-acetylation leads to the degradation of the protein (bioRxiv, doi:https://doi.org/10.1101/2020.11.22.393256). In this study, we were interested to investigate whether Api5 undergoes any other post translational modification(s) during normal physiological conditions or upon DNA damage prior to its degradation. Ubiquitination is a well-known post-translational modification that leads to degradation of the target protein. It is a complex process and requires multiple enzymatic functions in order to achieve degradation of its target proteins (7). Ubiquitin ligases (E3) are the most heterogeneous type of enzymes that carry out degradation by attaching ubiquitin moieties to the proteins (8). The multi-protein E3 ubiquitin ligase Skp, Cullin, F-box containing complex (SCF complex) helps in degradation of proteins that function during the cell cycle as well as in apoptosis (9). Most of the proteins that undergo ubiquitination also require phosphorylation as a secondary but necessary signal for their degradation. Since Api5 has earlier been reported to play a significant role in DNA damage induced apoptosis, it was important to investigate the mechanism of regulation of Api5 during DNA damage.

ATR is one of the sensor kinases that is involved in invoking a cell cycle arrest to allow for repair of damaged DNA (10). To perform its functions, ATR interacts with and phosphorylates BRCA1, CHK1, MCM2, RAD17, RPA2, SMC1 and p53/TP53 that collectively inhibit DNA replication as well as mitosis and promote DNA repair, recombination and apoptosis (11–17).

As earlier reports have shown Api5 to undergo post-translational degradation, it was pertinent to investigate the mechanism of its degradation during normal physiological conditions as well as during DNA damage. Our findings demonstrate Api5 to undergo poly-ubiquitination and thereby degrade via the proteasomal pathway during both normal physiological conditions as well as during DNA damage-induced apoptosis. We report FBXW2, a component of SCF (SKP1-Cullin1-F-box protein) E3 ubiquitin ligase to play a role during the proteasomal-mediated degradation of Api5. We also demonstrated that Api5 degradation upon DNA damage was regulated by ATR. Thus, we report ATR and FBXW2 as key regulators of Api5 function during physiological as well as during DNA damage conditions.

## Results

### Api5 undergoes ubiquitin proteasome-mediated degradation

To identify the potential pathway of Api5 degradation, cells were treated with different concentrations of chloroquine, a lysosomal pathway inhibitor and MG132, a proteasomal pathway inhibitor. Inhibition of the lysosomal pathway did not show any change in Api5 levels suggesting that Api5 degradation is not through the lysosomal pathway (Figure 1A and S1A). However, upon inhibition of the proteasomal pathway by varying the dose of MG132, showed increased levels of Api5 indicating that Api5 degradation is through the proteasomal pathway (Figure 1B and S1B). In order to obtain optimal accumulation of Api5, 10μM of MG132 was added to the cells and time-course experiments were performed. It was observed that inhibition of the proteasomal pathway for 4 hrs using 10 μM of MG132 showed optimal accumulation of Api5 (Figure 1C–D). After standardising the optimal dose and time for MG132 treatment, all further experiments were performed with that particular dose and Api5 levels were analysed. To further validate that the increase in protein level upon inhibition of the proteasome pathway was not as a result of increased protein synthesis, cells were treated with 10 μg/ml of cycloheximide, a protein synthesis inhibitor, for different time durations. Decrease in Api5 protein levels upon cycloheximide treatment confirmed the inhibition of Api5 protein translation (Figure 1E–F). To prove that the accumulation of Api5 is due to inhibition of proteasomal degradation and not due to increased protein synthesis, MCF7 cells were treated with a combination of cycloheximide and MG132. Accumulation of Api5 was observed upon treatment with MG132 even when protein synthesis was inhibited, thereby demonstrating that the degradation of Api5 is through the proteasomal pathway (Figure 1G–H). A plethora of studies have shown that most of the proteins that degrade via the proteasomal pathway undergo ubiquitination as a post-translational modification. Therefore, to verify whether Api5 undergoes ubiquitination, immuno-precipitation was performed using Api5 specific antibody. Immunoblotting of the Api5 immunoprecipitates revealed the presence of high mass ladders, which were further augmented following addition of MG132, indicating that Api5 undergoes poly-ubiquitination (Figure 1I).

**Figure 1.**
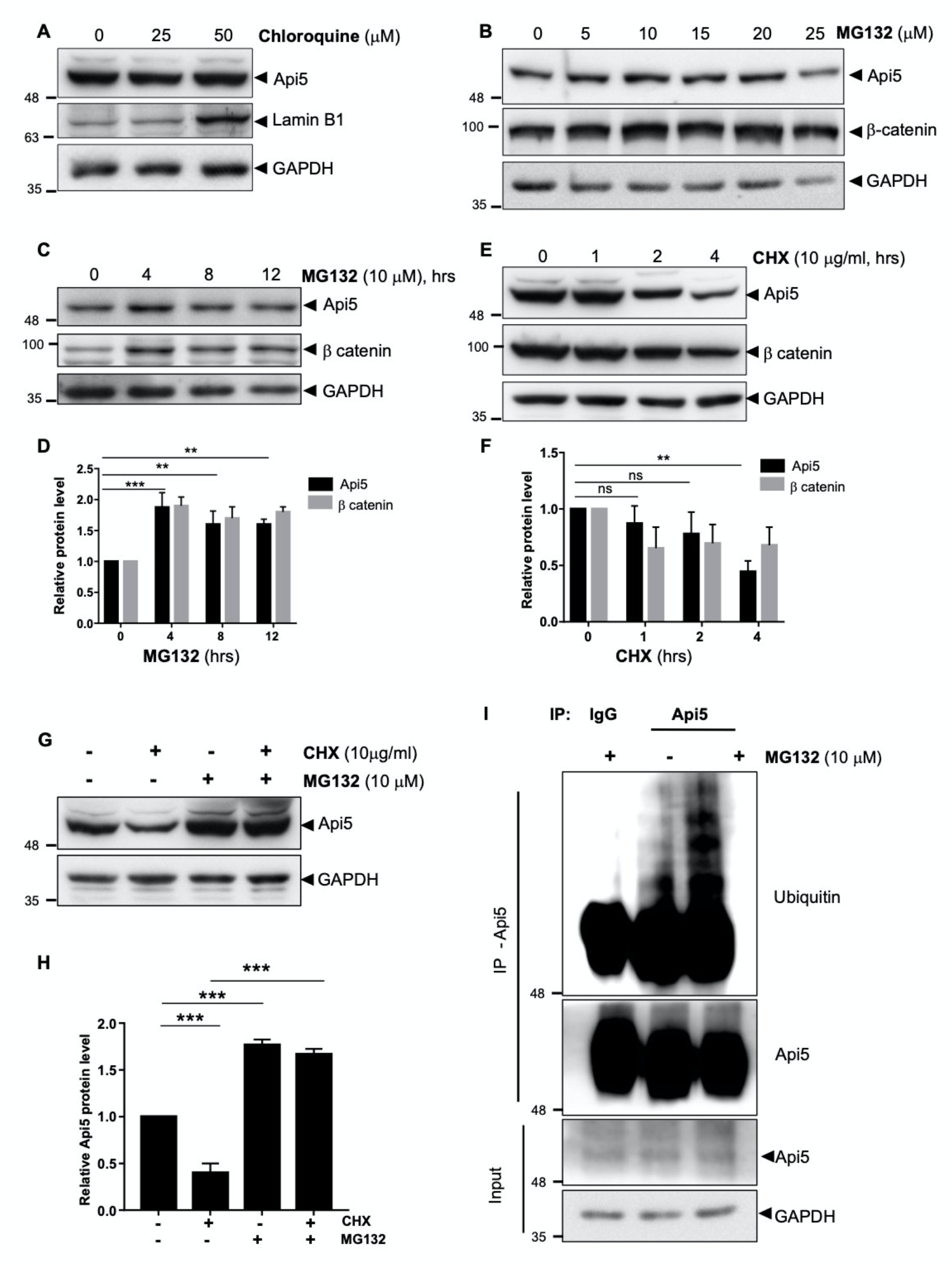
Api5 undergoes proteasomal degradation. (A) Western blot analysis showing protein levels of Api5 in MCF7 cells upon treatment with different doses of chloroquine. Lamin B1 was used as a positive control and GAPDH was used as a loading control. MCF7 cells treated with (B) varying doses of MG132 for 8hrs and (C) 10 μM of MG132 for different time points. (D) Api5 levels were quantified and normalised to GAPDH. (E) Api5 levels were analysed in MCF7 cells treated with cyclohexamide for different time points and (F) quantification afer normalising to GAPDH. β catenin was used as the postive control for MG132 and cyclohexamide treatment. (G) MCF7 cells were treated with 10μg/ml of cycloheximide and 10μM MG-132 and Api5 levels were analysed using immunoblotting. (H) Api5 levels were quantified and normalised to GAPDH. (I) MCF7 cells treated with MG132 were lysed and immunoprecipitation was performed using Api5 specific antibody and immunoblotting using ubiquitin antibody.

### F-box protein FBXW2 degrades Api5

F-box proteins are the substrate receptor component of SCF E3 ubiquitin ligases and they promote poly-ubiquitination-mediated degradation of their target proteins. To investigate the F-Box proteins involved in the degradation of Api5, cells were transfected with plasmids containing different F-box genes individually. Levels of Api5 were analysed following ectopic expression of myc- and flag-tagged FBOX proteins by immunoblotting. Of the 56 F-Box proteins screened, Api5 levels were significantly decreased following ectopic expression of FBXL6, FBXL8, FBXW2, FBXW4, FBXW9, FBXO3 and FBXO6 (Figure 2A). Quantification of Api5 levels revealed that the ectopic expression of FBXW2, FBXO3, and FBXO6 resulted in reduction of Api5 to its minimal level (Figure 2B).

**Figure 2.**
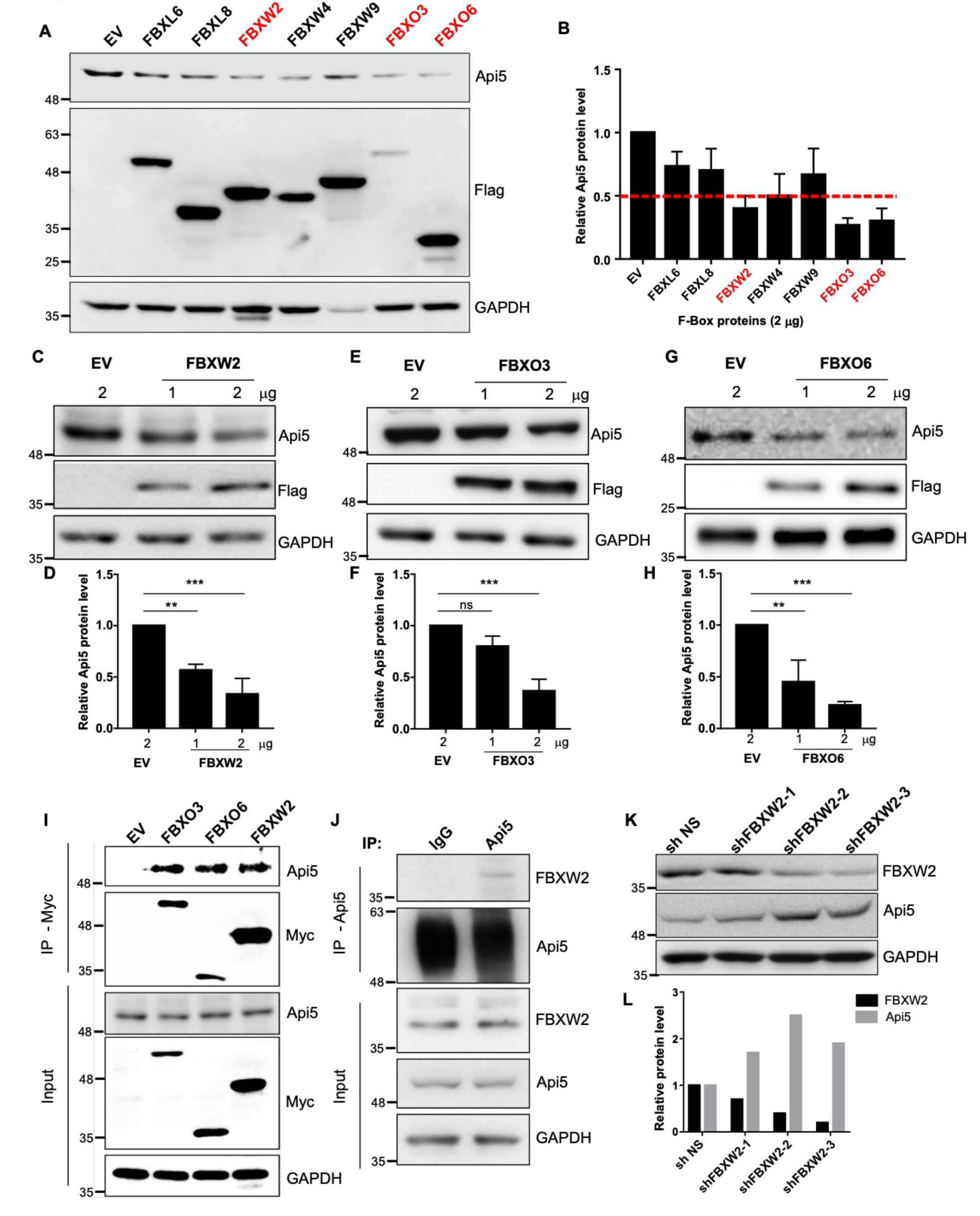
FBXW2 degrades Api5. Plasmids containing various flag-tagged F-box genes were ectopically expressed in MCF7 cells by transient transfection. These cells were lysed and Api5 levels were analysed using (A) immunoblotting and (B) quantified after normalising with GAPDH. Ectopic expression of F-box proteins was determined by immunoblotting with Flag. antibody MCF7 cells transfected with 1 and 2 μg of (C) FBXW2, (E) FBXO3 and (G) FBXO6 and Api5 levels were analysed using immunoblotting and (D), (F), (H) quantified after normalising with GAPDH. Empty vector (EV; 2μg) transfected cells lysates were used as control. (I) MCF7 cells transfected with myc-tagged FBXW2, FBXO3 and FBXO6 containing plasmids were lysed and immunoprecipitatied using myc antibody. Api5 levels were analysed by immunoblotting with Api5 specific antibody. (J) Endogenous Api5 was pulled down in MCF7 cells and FBXW2 interaction was analysed using immunoblotting. (K) HeLa cells stably expressing various shRNA constructs of FBXW2 were lysed and Api5 and FBXW2 levels were analysed using immunoblotting. (L) FBXW2 and Api5 levels were quantified and normalised to GAPDH.

To demonstrate a dose dependent degradation of Api5 by the selected F-Box proteins, cells were transfected with either vector control or plasmids containing FBXW2, FBXO3 and FBXO6 genes, and Api5 protein levels were analysed. It was observed that FBXW2, FBXO3 and FBXO6 degrade Api5 in a dose-dependent manner (Figure 2C–H). To prove that F-Box protein-mediated degradation of Api5 is through the proteasomal pathway, cells were treated with MG132 upon overexpression of FBXW2, FBXO3 and FBXO6. Api5 levels were restored upon MG132 treatment in the transfected cells suggesting that F-Box proteins degrade Api5 through the proteasomal pathway (Figure S2A-F).

In order for Api5 to undergo ubiquitin-mediated proteasomal degradation, the selected F-Box proteins must interact with Api5. To examine this, myc-tagged FBXW2, FBXO3 and FBXO6 were ectopically expressed and immuno-precipitation was carried out using anti-myc antibody. Presence of Api5 in the immune-precipitates of F-box proteins indicate that FBXO3, FBXO6 and FBXW2 interact with Api5 (Figure 2I).

Among the selected F-box proteins, FBXW2 has been reported to act as a tumour suppressor in lung carcinomas by degrading oncogenic SKP2 (18). Moreover there are multiple studies that suggest Api5 can act as a potential oncogene. We therefore speculated that FBXW2 may be potential candidate F-box protein to inhibit the anti-apoptotic function of Api5 and therefore we carried detailed studies to understand the mechanism of its regulation. First we examined the interaction of FBXW2 and Api5 at the endogenous level by co-immunoprecipitation assay. Immunoblotting results showed the presence of FBXW2 in the immuno-precipitates of Api5, indicating that Api5 and FBXW2 interact at the endogenous level (Figure 2J).

Next, we examined whether FBXW2 controlled basal levels of Api5. To address this, FBXW2 was depleted in cells using three independent shRNAs and a non-specific shRNA was used as control. Immunoblotting results revealed that expression levels of Api5 increased following depletion of FBXW2 (Figure 2K–L). Taken together, it may be concluded that FBXW2 plays a critical role in restricting the cellular levels of Api5 by directing the cells for proteasomal degradation.

### Api5 protein level reduces during DNA damage-induced apoptosis

To investigate the regulation of Api5 upon DNA damage, cells were treated with different concentrations of chemical and UV-radiation damage. Upon treatment with different doses of camptothecin (CPT) and etoposide (ETP), DNA damage was induced that was confirmed by observing the activation of Chk1 and Chk2 using immunoblotting as well as the presence of phosphorylated RPA foci by immuno-fluorescence (Figure S3A-D). Similarly, UV-irradiation led to the phosphorylation of Chk1 and the presence of pRPA foci, thus confirming DNA damage in the cells (Figure S3E-F). To demonstrate that DNA damage caused by chemicals as well as UV-irradiation can initiate apoptosis in the cells, apoptotic markers were analysed where increased levels of cleaved PARP and caspase 9 were observed (Figure S3 G-I). After standardising the dose and time for each DNA damaging agent, the levels of Api5 were analysed in the treated cells. It was observed that cells treated with camptothecin, etoposide and UV-irradiation showed reduced levels of Api5 (Figure 3A–F). This reduction in Api5 levels may be due to decrease in transcription of Api5. However, RT-PCR data demonstrated unaltered Api5 transcript levels upon DNA damage (Figure 3G–L), thereby indicating that Api5 may be regulated at the post-transcriptional level following DNA damage.

**Figure 3.**
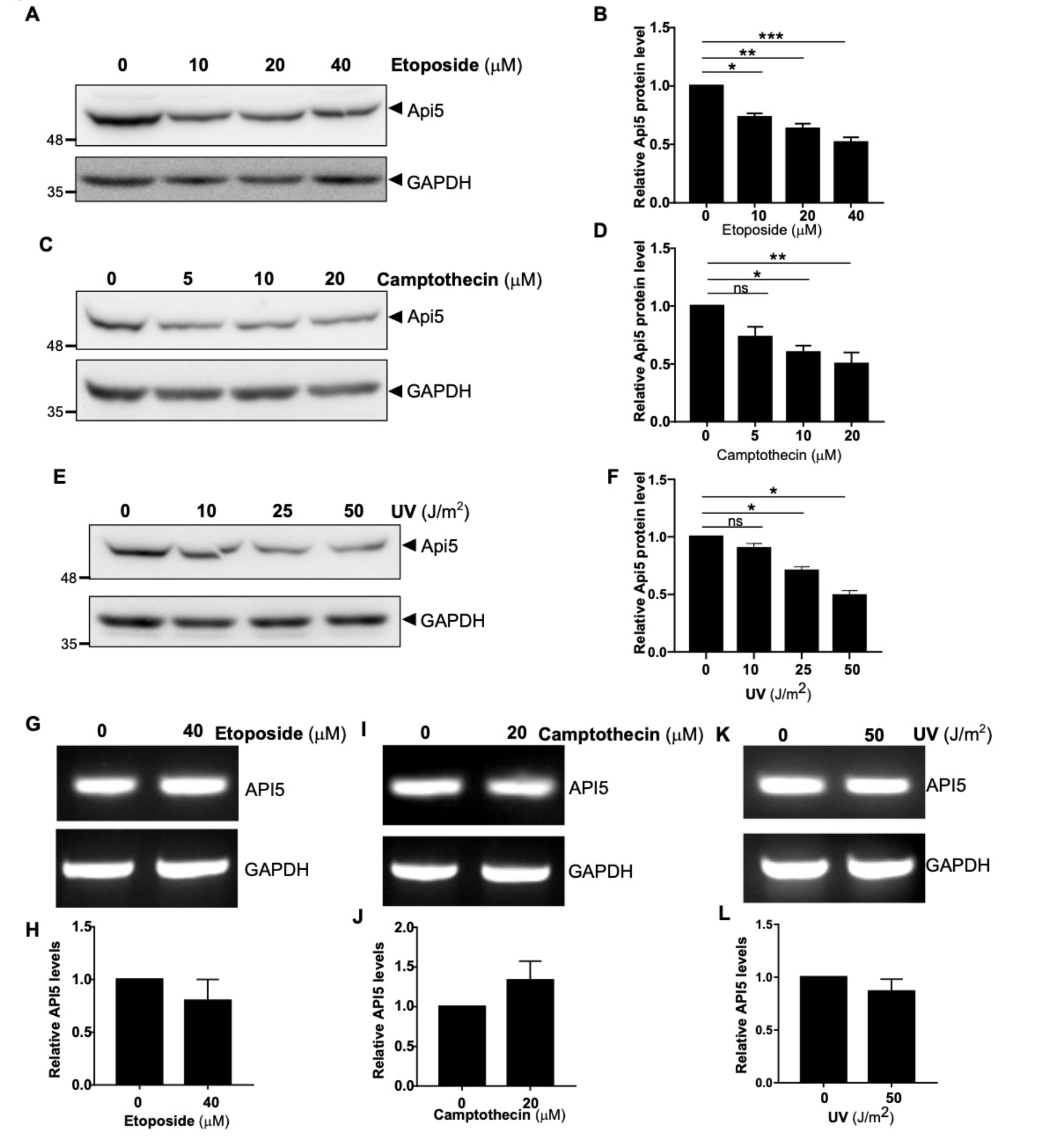
Api5 undergoes post-translational degradation upon DNA damage. MCF7 cells treated with varying doses of (A) etoposide for 24 hrs, (C) camptothecin for 16hrs and (E) UV for 12 hrs were lysed and Api5 levels were analysed using immunoblotting and quantified after normalising with GAPDH in (B, D, and F). API5 transcript levels upon (G) etoposide, (I) camptothecin and (K) UV treatment were analysed using semi-quantitative PCR and (H, J, L) quantified after normalising to GAPDH.

### Api5 undergoes DNA damage-mediated degradation through ubiquitin-proteasomal pathway by FBXW2

Previous experiments suggested that Api5 undergoes degradation via the proteasomal pathway during physiological conditions and Api5 levels reduce upon DNA damage. It was hypothesised that Api5 undergoes proteasomal mediated degradation upon DNA damage. As proteasome complex is localised in the cytoplasm while Api5 is a nuclear protein, we first investigated the localisation of Api5 following UV damage. In control cells, Api5 was observed to be present in the nucleus while upon UV treatment, most cells showed nuclear as well as cytoplasmic localisation of Api5 (Figure 4A–B). Interestingly, the increase in percentage of cells showing only cytoplasmic presence of Api5 upon UV-irradiation in the 8 and 12 hrs post treatment, suggest that Api5 compartmentalisation changes upon UV damage (Figure 4B).

**Figure 4.**
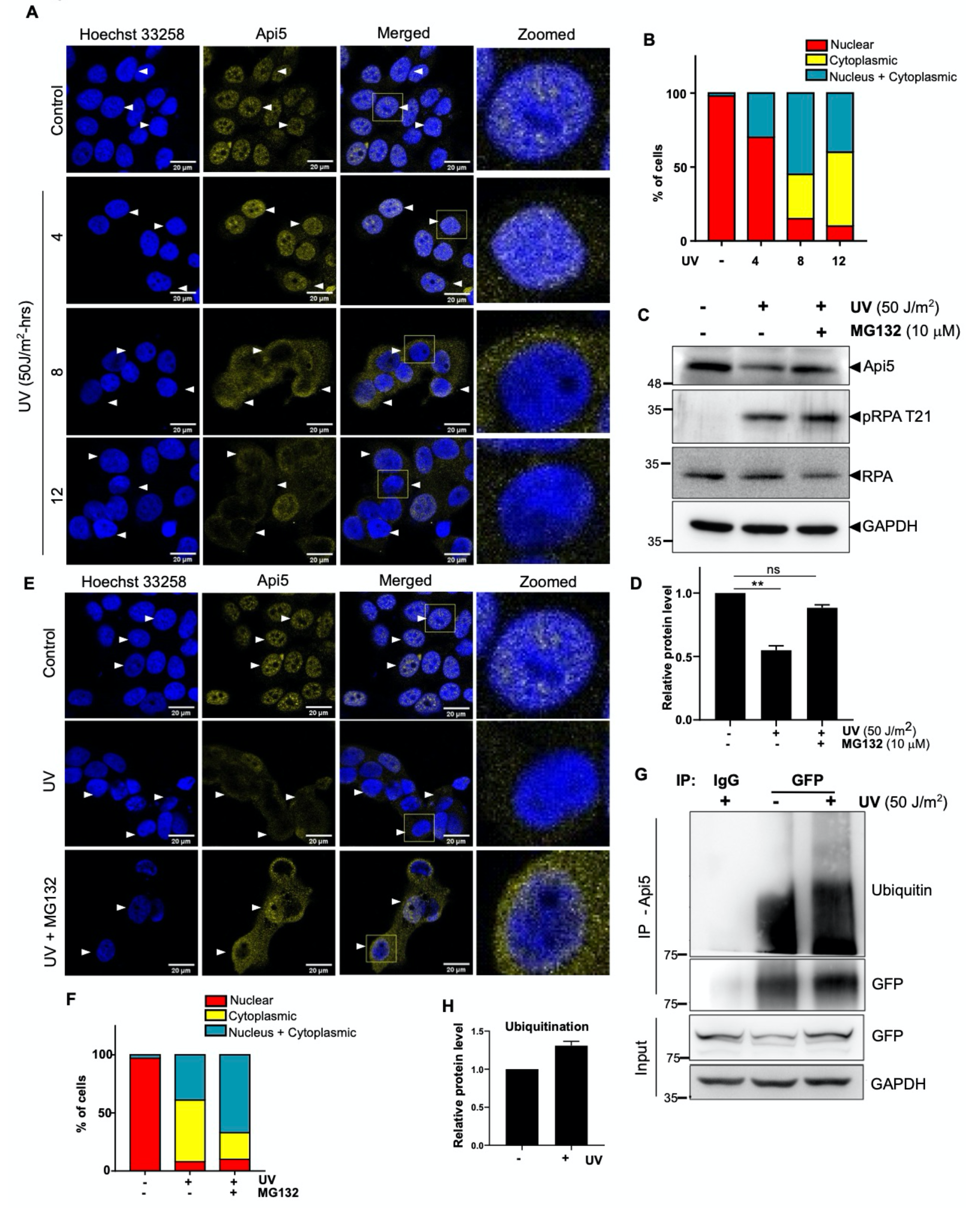
Api5 undergoes proteasomal pathway mediated degradation upon DNA damage. (A) MCF7 cells treated with 50 J/m^2^ UV and fixed post 4, 8 and 12 hrs of irradiation. Immunofourescence was performed using Api5 specific antibody and imaged in Leica SP8 microscope at 63X oil immersion objective (scale bar: 20 μm) and (B) cells with different Api5 staining pattern were manually counted and analysed. (C) MCF7 cells treated with 50 J/m^2^ UV and 10 μM MG132 for 4hrs were lysed after 12 hrs post UV treatment and Api5 levels were analysed using western blotting. (D) The fold change of Api5 was quantified after normalisation with GAPDH. (E) Immunofluorescence was performed on MCF7 cells irradiated with 50 J/m^2^ UV and with and without MG132. (F) Api5 localisation changes were manually analysed. (G) mCherry-tagged Api5 over-expressing MCF7 cells treated with 50 J/m^2^ UV and 10 μM MG132 for 4 hrs were lysed 12 hrs post UV tretment and immunoprecipitation was performed using GFP specific antibody. Western blotting analysis was performed to analyse the ubiquitination status of Api5 using ubiquitin specific antibody. (H) Ubiquitin levels of GFP pull down lysates were quantified after normalising to GFP and represented as bar graph.

To investigate whether Api5 undergoes proteasomal-mediated degradation upon UV damage, Api5 levels were analysed upon inhibition of the proteasomal pathway by MG132 treatment following UV damage. DNA damage was confirmed by the presence of activated pRPA32 by immunoblotting. Restored levels of Api5 were observed upon inhibition of the proteasomal pathway following UV damage suggesting that DNA damage-mediated degradation of Api5 was also through the proteasomal pathway (Figure 4C–F). To illustrate whether ubiquitination of Api5 precedes its degradation via the proteasomal pathway, mCherry-tagged Api5 overexpressing stable cells were immunoprecipitated using GFP-specific antibody. It was observed that Api5 undergoes poly-ubiquitination upon DNA damage (Figure 4G–H). These results overall suggested that Api5 undergoes proteasome pathway mediated degradation upon DNA damage.

### ATR regulates localisation and degradation of Api5 post DNA damage

Preceding results demonstrated that Api5 undergoes ubiquitin-proteasomal mediated degradation upon DNA damage. ATR kinase is one of central kinases responsible for relaying DNA damage response and repair. It phosphorylates numerous proteins associated with the DNA damage response and repair process. We investigated the role of ATR in proteasomal degradation of Api5 during DNA damage. Immunoblotting results showed that UV-induced reduction of Api5 was blocked following inhibition of ATR (Figure 5A–B). These results were further corroborated using immunofluorescence studies where the cytoplasmic localisation of Api5 upon UV damage decreased upon inhibition of ATR (Figure 5C–D). Interestingly Api5 was observed to form speckles inside the nucleus upon blocking the ATR kinase activity in UV-treated cells. These data prove that ATR regulates DNA damage mediated degradation of Api5. Immunoprecipitation studies were performed to examine the interaction between ATR and Api5 in cells stably expressing mCherry-tagged Api5 following UV damage. Results demonstrated the presence of ATR in the immunoprecipitates of Api5 (Figure 5E). This interaction was further confirmed by a reciprocal immunoprecipitation study (Figure 5F). Presence of Api5 in the ATR immunoprecipitates confirmed the interaction between Api5 and ATR.

**Figure 5.**
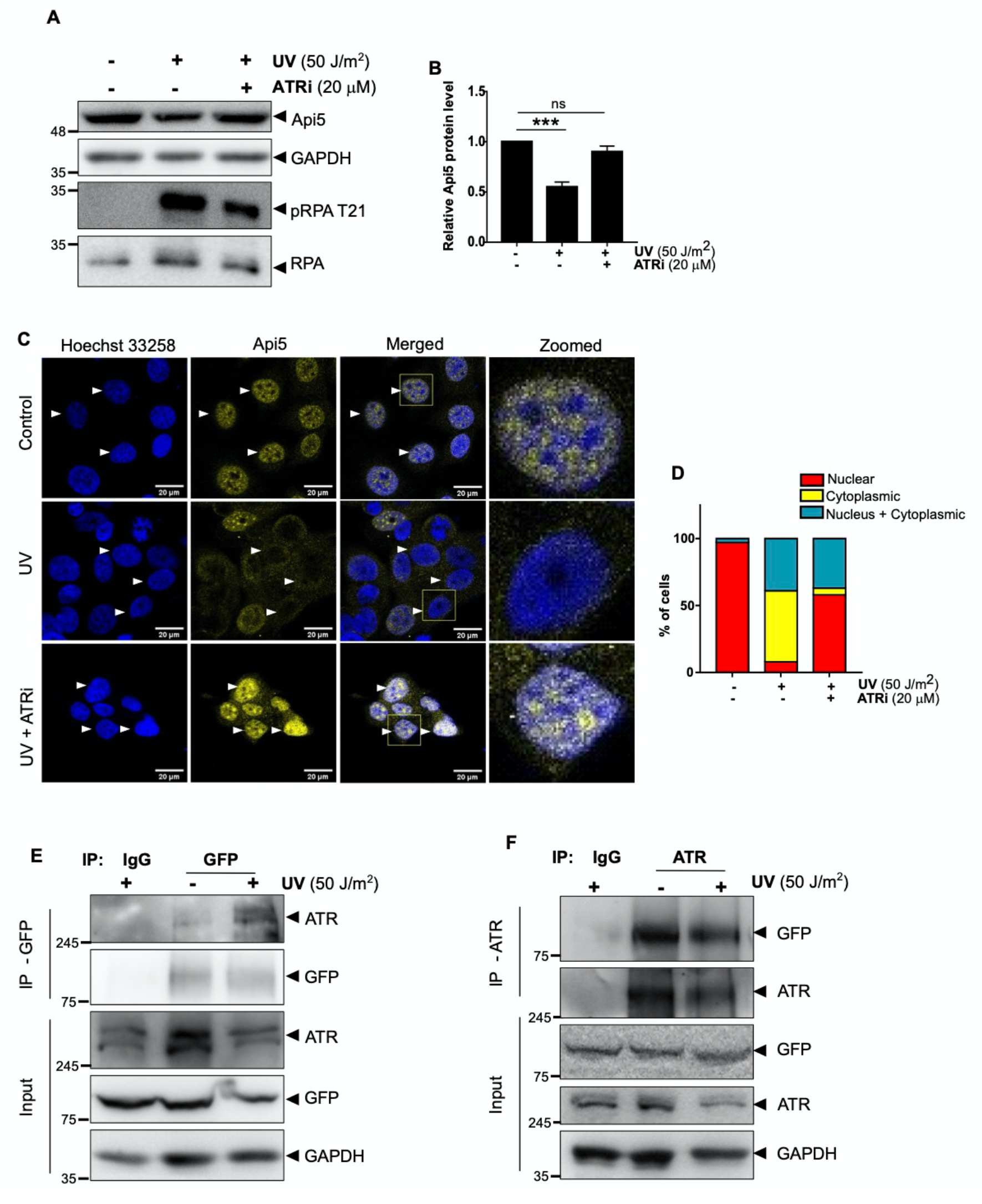
ATR interacts and regulates Api5 upon UV damage. (A) MCF7 cells were treated with 50 J/m^2^ UV and ATRi (20μM VE821 for 12 hrs) were lysed 12 hrs post UV and levels of Api5, pRPA and RPA were analysed using western blotting. (B) The fold change of Api5 was quantified after normalisation with GAPDH. (C) Immunofluorescence assay was performed on MCF7 cells treated with 50 J/m^2^ UV and ATRi (20μM VE821 for 12 hrs) using Api5 specific antibody and (D) Api5 localisation was manually analysed. mCherry-tagged Api5 over-expressing MCF7 cells treated with 50 J/m^2^ UV were lysed 12 hrs post treatment and immunoprecipitation was performed using (E) GFP and (F) ATR-specific antibody. Western blotting analysis was performed to analyse the presence of ATR and GFP respectively.

### ATR phosphorylates Api5 at serine 138 and regulates apoptosis upon DNA damage

Earlier studies have demonstrated that ATR/ATM phosphorylates their substrates at SQ/TQ sites. As ATR was found to be an interactor and regulator of Api5 upon DNA damage, we went ahead to investigate whether ATR can phosphorylate Api5 upon DNA damage. Analysis of Api5 amino acid sequence in Han et al., (2) showed the presence of only two (SQ/TQ) sites on Api5 where ATR could potentially phosphorylate Api5. One at serine 138 and the other at threonine 108 amino acid residues (Figure 6A). To assess whether ATR could phosphorylate Api5, mCherry-tagged Api5 stable cells were lysed and Api5 was immunoprecipitated with anti-GFP antibody and immunoblotting was performed using phospho-serine specific antibody. Presence of phospho-serine at the size of Api5 confirmed that Api5 undergoes phosphorylation upon DNA damage (Figure 6B–C). This phosphorylation could be a signal for Api5 to undergo ubiquitin-mediated proteasomal degradation. To understand the role of this phosphorylation on the stability of Api5, site-directed mutagenesis was performed to mutate serine 138 and threonine 108 to alanine. mVenus tagged wild-type, S138A and T108A mutants of Api5 were ectopically expressed and their stability were analysed upon DNA damage by immunoblotting. It was observed that upon DNA damage, protein levels of wild type and T108A Api5 reduced while Api5 S138A remained unaltered suggesting that phosphorylation at serine 138 of Api5 is responsible for its degradation (Figure 6D–E). In order to understand the function of Api5 S138A mutant upon UV damage, protein expression of various DNA damage repair and apoptotic proteins were analysed in cells expressing wild type and S138A Api5 mutant. Levels of pro-apoptotic proteins like cleaved PARP and caspase 9 were found to be elevated in wild-type cells upon UV damage while no change in protein levels of cleaved PARP or caspase 9 were observed in cells expressing Api5 S138A mutant, suggesting that ATR-mediated degradation of Api5 plays a significant role in the regulation of DNA damage induced apoptosis (Figure 6F–H). The reduction in the protein expression of apoptotic markers in Api5 S138A mutant cells may be due to reduced sensing of UV damage. It was therefore interesting to look for other DNA damage markers. As expected there was a significant difference in levels of total p53 and activation of p53 at serines 15, 37 and 46 upon DNA damage. Cells expressing Api5 S138A mutant showed reduced activation of p-p53 S15, p-p53 S37, p-p53 S46 as well as total p53 when compared to cells expressing wild-type Api5 upon UV damage (Figure 6I–M), suggesting that phosphorylation of Api5 at serine 138 not only regulates the stability of the protein but also desensitises cells towards DNA damage as well as DNA damage induced apoptosis.

**Figure 6.**
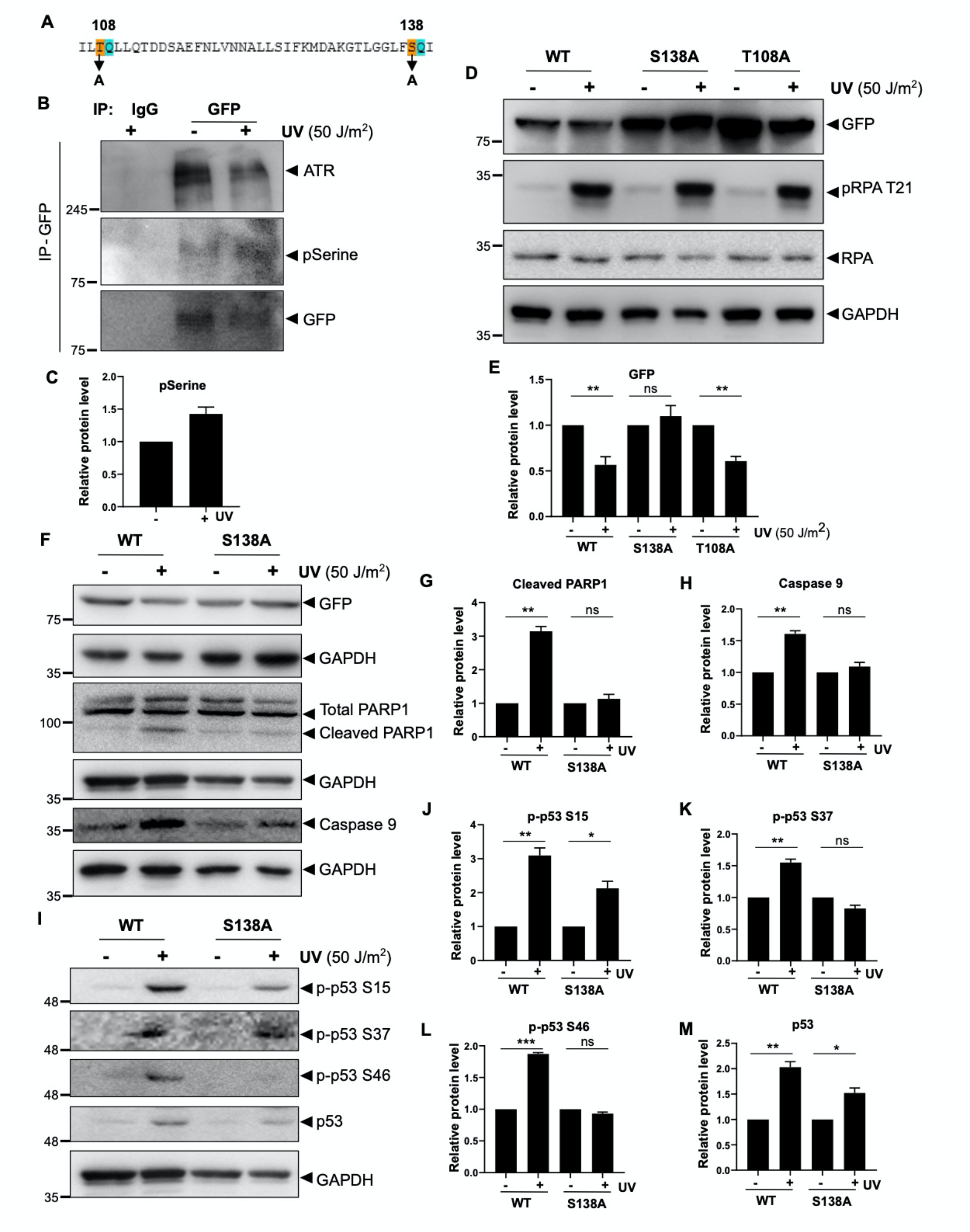
ATR phosphorylates Api5 at S138 to regulate DNA damage response. (A) Sequence showing the potential ATR phosporylation sites in Api5 protein highlighted at T108 and S138 residues. (B) mCherry-tagged Api5 over-expressing MCF7 cells treated with 50 J/m^2^ UV were lysed 12 hrs post treatment and immunoprecipitation was performed using GFP-specific antibody. Western blotting analysis was performed to analyse the presence of phospho-serine and ATR. (C) pSerine levels of GFP pulldown lysates were quantified after normalising to GFP and represented as bar graphs. Serine at 138 and threonine at 108 residues on Api5 were mutated to alanine by site-directed mutageneis. MCF7 cells transfected with mVenus tagged-WT, S138A and T108A-Api5 mutants were UV irradiated (50 J/m^2^) post 24hrs of transfection and lysed post 12 hrs of UV treatment. Western blotting was performed to check the levels of (E) Api5, (F) PARP and caspase 9, (I) p53-S15, p53-S37, p53-S46 and total p53 and (E, G, H, J, K, L and M) quantified after normalising to the loading control GAPDH.

### De-acetylated form of Api5 undergoes degradation during DNA damage

Studies from our laboratory have demonstrated that p300 and HDAC1 are the enzymes involved in the regulation of the stability of Api5. p300 acetylates Api5 at lysine 251 (K251) and maintain its stability while the HDAC1 deacetylates and leads to the degradation of Api5 (bioRxiv; doi: https://doi.org/10.1101/2020.11.22.393256). As our current data also reports Api5 to undergo ubiquitin-proteasome pathway mediated post translational degradation, it was interesting to investigate the role of p300-mediated acetylation and HDAC1 deacetylation on the regulation of stability and anti-apoptotic functions of Api5 following DNA damage. UV-irradiated cells were treated with romidepsin, a small chemical inhibitor of HDAC1 and levels of Api5 were analysed post UV damage. Api5 levels remained unaltered upon inhibition of HDAC1 in UV-irradiated cells suggesting that DNA damage induced degradation of Api5 requires de-acetylation by HDAC1 (Figure 7A–B).To check for other molecular players involved in the acetylation and stability of Api5 upon UV damage, cells were treated with UV-irradiation and protein levels of p300 and HDAC1 were analysed. p300 levels were found to be reduced upon UV damage while HDAC1 levels remained unaffected (Figure S4A-F). To demonstrate that acetylation of Api5 at K251 is required for maintaining stability of Api5 following DNA damage, mVenus-tagged K251 mutant of Api5, K251Q which is a constitutive acetylation mimic mutant was ectopically expressed along with wild-type Api5 followed by UV-irradiation and the levels of Api5 were analysed post DNA damage. Wild type Api5 levels were observed to reduce while the K251Q mutant Api5 protein levels did not alter upon UV-irradiation suggesting that acetylation at lysine 251 plays a crucial role in maintaining the stability of Api5 (Figure 7C–D). Stability of proteins is generally considered to be a direct evidence for their functions, therefore the protein expression of various DNA damage and apoptotic markers were analysed in Api5 wild-type and K251Q mutant expressing cells upon UV damage. Wild-type Api5 showed increased levels of cleaved PARP while the other Api5 acetylation mutants failed to show any difference in the protein expression levels (Figure 7C–E). To understand the role of Api5 and K251 acetylation of Api5 on cell survival upon DNA damage, wild type and K251 mutants were stably expressed in Api5 knocked down stable cells. Cells with Api5 knock down showed reduced survival upon camptothecin and etoposide treatment compared to the control cells while upon rescue with wild type Api5, cell survival was restored (Figure 7 F–G). K251Q mutant of Api5 also facilitated cell survival similar to wild type Api5, however K251A mutant of Api5 was not able to rescue the cell survival upon DNA damage. These results suggest that Api5 acetylation at K251 plays a significant role during regulation of DNA damage-induced apoptosis.

**Figure 7-.**
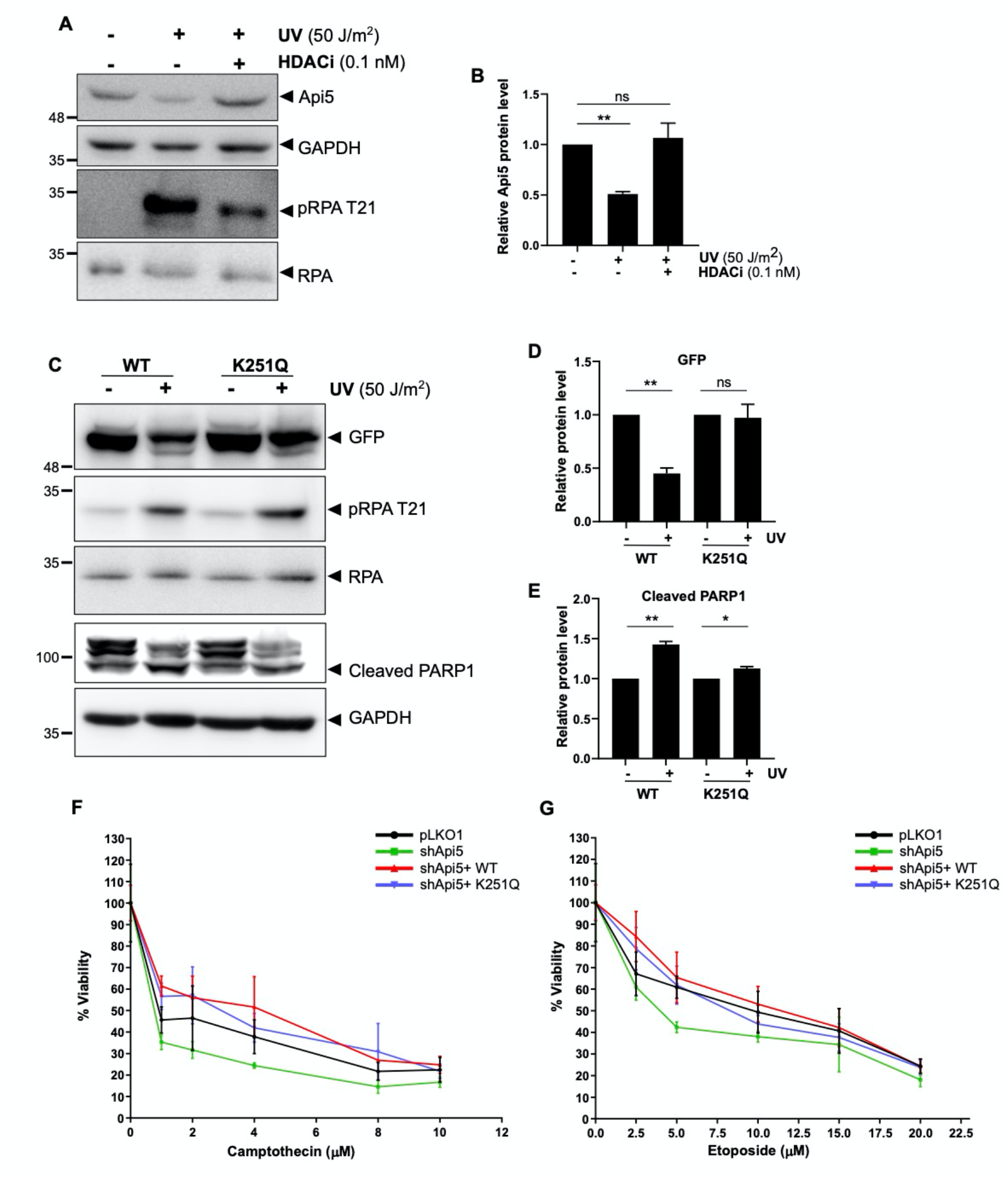
Acetylation of Api5 at K251 regulates DNA damage induced apoptosis. (A) MCF7 cells treated with HDACi (0.01nM romidepsin for 12hrs) and UV (50 J/m^2^) were lysed post 4 hrs of UV irradiation and Api5 levels were analysed using western blotting. (B) The fold change of Api5 was quantified after normalising to GAPDH. MCF7 cells were transfected with mVenus-tagged wild type and K251Q mutant constructs and treated with 50J/m^2^ UV post 24 hrs of transfection and lysed 12 hrs post UV irradiation. Western blotting was performed to check for levels of (C) GFP, PARP. (D and E) quantification of the western blots after normalising to the loading control GAPDH. Viability of MCF7 cells stably expressing shApi5 and K251 mutants upon treatment with varying concentrations of (F) camptothecin and (G) etoposide for 24 hrs were measured using MTT assay.

## Discussion

Api5 is an anti-apoptotic protein that helps in cell survival during serum starvation conditions (1). Later it was found that the anti-apoptotic function of Api5 was not only confined to serum starvation conditions but it also could potentially inhibit apoptosis during infectious and DNA damage conditions (19) (3) (6). However, the mechanism by which Api5 is able to regulate itself during stress conditions like serum starvation, infections and DNA damage is yet to be understood. In this study we report that Api5 undergoes degradation upon DNA damage induced apoptosis. UV-irradiation and chemical damage resulted in reduction in Api5 protein levels. Our results suggest that post translational degradation of Api5 upon DNA damage induced apoptosis is through the proteasomal pathway. Ubiquitination is the post translational modification that targets Api5 towards the proteasome machinery. However, we are not dismissing the possibility that Api5 may be regulated via other post-translational modifications and/or other pathways involved in the post-translational degradation of Api5.

FBXW2 is a SCF-type of E3 ubiquitin ligase that is involved in the ubiquitin-mediated degradation of a potential cell cycle regulator SKP2 (18). It has already been reported that in lung carcinomas, SKP2, a potential oncogenic protein gets degraded by FBXW2 which thus acts as a tumor suppressor. Api5 is also found to be upregulated in different types of cancers like cervical, oesophageal and lung cancers (18,20–22). Multiple studies by various groups suggest that Api5 can also be a potential oncogenic factor which upon its down regulation could make cells vulnerable towards DNA damage mediated cell death (3). Our studies confirmed that FBXW2 interacts with Api5 and promotes its degradation through ubiquitination during physiological conditions as well as during DNA damage induced apoptosis.

Phosphorylation is a secondary but an essential signal for degradation of multiple proteins via the proteasomal pathway. ATR is one of the main sensor kinase that upon sensing DNA damage, activates a cell cycle checkpoint response by phosphorylating specific proteins like Chk1 and p53 (10,12,13). Our results indicate that ATR binds to Api5 and phosphorylates it at the serine 138 residue. Phosphorylation of Api5 at serine138 is required for its degradation during DNA damage induced apoptosis. Upon DNA damage Api5 first gets de-acetylated by HDAC1. This de-acetylated Api5 undergoes post-translational degradation through the proteasomal pathway after getting phosphorylated by ATR. The phosphorylated form of Api5 localises to the cytoplasm and undergoes degradation post-DNA damage while the nuclear localised form of Api5 is acetylated and stabilised by p300 histone acetyltransferase. The acetylated form of Api5 that does not undergo degradation upon DNA damage may be playing a role in DNA repair and/or differential regulation of apoptosis upon DNA damage. In our studies we identified ATR as a major regulator of Api5 however we are also aware that other regulators like ATM and DNA-PK may also play a role that is yet to be investigated. Increased levels of Api5 are found to be associated with higher aggressiveness of cancers and poor cancer patient survival (23). Ectopic expression of Api5 can also promote properties like invasion, migration and metastasis in cancers (24). Depletion of Api5 induces apoptosis while over-expression of Api5 provides chemo-resistance to certain cancers (23). These studies demonstrate Api5 to be a potential oncogene and it is essential to understand the mechanism of regulation of stability as well as function of Api5 upon DNA damage. By understanding the molecular mechanism of regulation of Api5, multiple cancers can be targeted using techniques or molecules which specifically block the function of Api5. These molecules can be used as potential therapeutic agents to treat cancers where Api5 is over-expressed. Taken together we report FBXW2 and ATR as the novel regulators of Api5 upon DNA damage. Upon DNA damage HDAC1 de-acetylates Api5 and this de-acetylated Api5 undergoes post-translational degradation. Upon UV irradiation, ATR phosphorylates Api5 at S138 residue. This signals FBXW2 E3 ubiquitin ligase to target the protein for ubiquitination and degradation as summarised in Figure 8.

**Figure 8:**
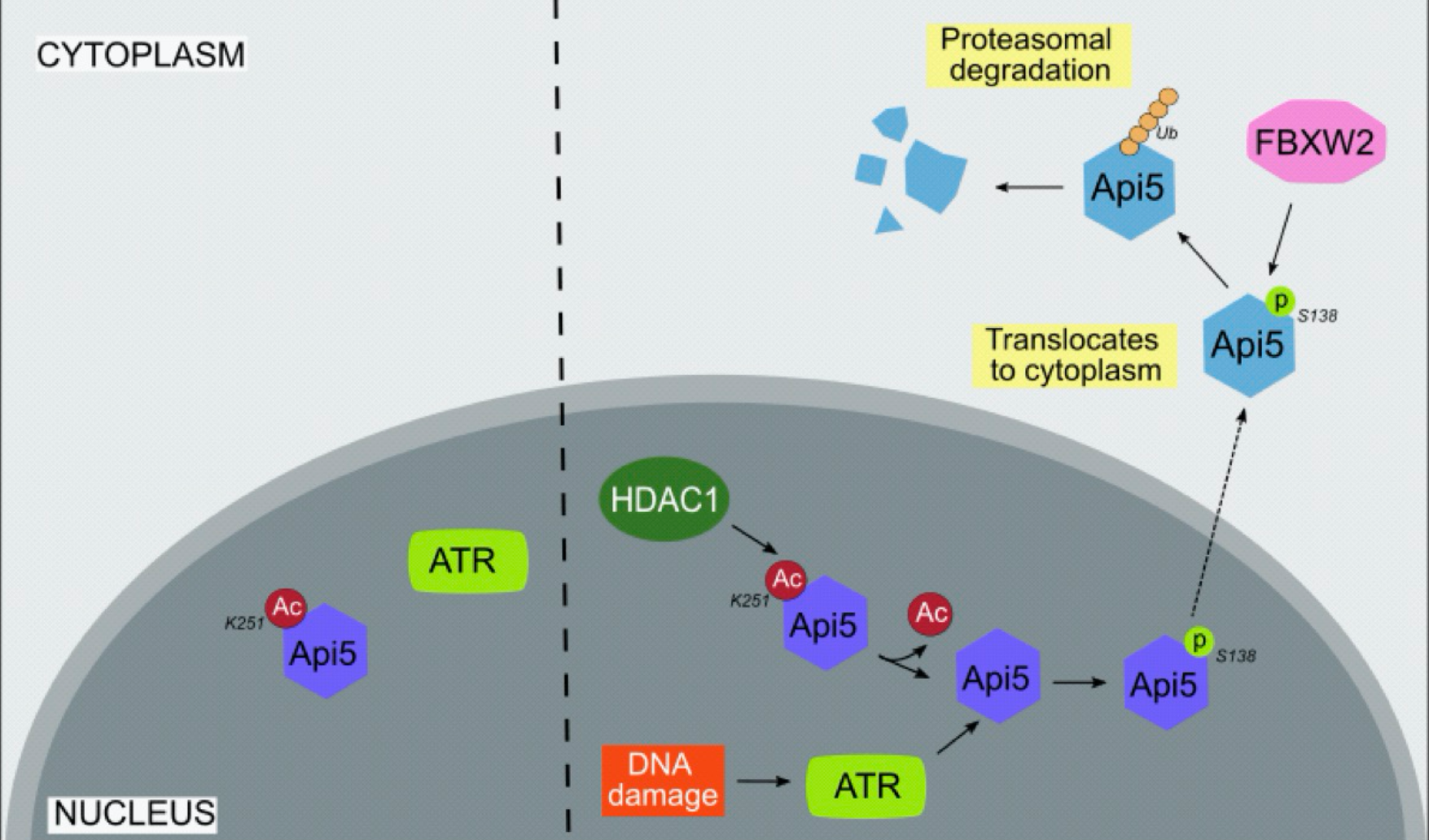
Schematic representation summarising the regulation of Api5 upon DNA damage. A proposed model summarising the pathway of regulation of Api5 by its key regulators HDAC1, ATR and FBXW2 upon DNA damage.

## Materials and methods

### Chemicals and antibodies

Camptothecin (C9911), Etoposide (E1383), Protease inhibitor cocktail (P2714), dimethyl sulfoxide (DMSO), propidium iodide (P4170) Chloroquine (C6628), MG132 (M7449), Cyclohexamide (C1988), polybrene (Hexadimethrine bromide; H9268), thiazolyl blue tetrazolium (MTT; M5655) and dimethyl sulfoxide (DMSO; D8418) were purchased from Sigma-Aldrich; romidepsin (FK228, Depsipeptide) was from Seleck Chemicals, USA and the selective ATR kinase ATR inhibitor (VE 821) was purchased from Axon Medchem, USA. Lipofectamine 2000 (H668), Alexa flour 488 goat anti-rabbit (A11034) and Phalloidin 568 (A12380) were purchased from Thermo Fisher Scientific, USA. FBXW2 (ab 83467), GFP (ab290), p300 (ab10485), ATR (ab2905), RPA T21 (ab61065), Caspase 9 (ab2324) antibodies were obtained from Abcam, USA. HDAC1 (sc-8410), Chk1 (G4, sc-8408), Ubiquitin (sc-8017) were purchased from Santacruz while Api5 (PAB7951) was from Abnova, Taiwan. Chk1 S345 (2348), Chk2 T68 (2661), Chk2 (3440), p53 S15 (9284), p53 S37 (9289), p53 S46 (2521), RPA 32 (2208) and c-Myc (9402) were from Cell Signaling Technology, USA while p53 (247A) and GAPDH (G9545) were from Bethyl Laboratories, USA and Sigma-Aldrich respectively. Hoechst 33258 (H3569) was purchased from Molecular Probes, Thermo Fisher Scientific, USA.

### Plasmids

CSII-EF-MCS plasmid was a gift from Dr. Sourav Banerjee (NBRC, Manesar, India). pCAG-HIVgp and pCMV-VSV-G-RSV-Rev plasmids were purchased from RIKEN BioResource Centre, Japan. pLKO.1 eGFP was a kind gift from Dr. Sorab Dalal, ACTREC, India. shApi5 sequence was cloned into the same using Addgene pLKO.1 protocol. Lentiviral pGIPZ shFBXW2 plasmids were a kind gift from Prof. Michael R. Green, University of Massachusetts Medical School, USA. psPAX2 and pMD2.G were obtained from Dr. Didier Trono (Addgene, Watertown, MA, USA). Human cDNA encoding FBOX genes were procured from Origene (Rockville, MA, USA). mVenusC1 was gifted by Dr. Jennifer Lippincott-Schwartz, NIH, USA in which Api5 was cloned. Api5 was made siRNA-resistant and K251 and S138A mutants were generated using site directed mutagenesis.

### Cell culture and drug treatments

MCF7 cells were obtained from the European Collection of Cell Cultures (ECACC). HEK293T and HeLa cells were a gift from Dr. Jomon Joseph (NCCS, Pune) and Dr. Sorab Dalal respectively. The cells were maintained in 100 mm dishes (Thermo Fisher Scientific, USA, 172931) and were grown in high-glucose Dulbecco’s Modified Eagle Medium (DMEM; Lonza Group AG, Switzerland, 12-604F) containing 10% heat inactivated FBS (Thermo Fisher Scientific, USA, 10270106) and 100 units/mL penicillin-streptomycin (Lonza Group AG, Switzerland, 17-602E) and incubated at 37°C in humidified 5% CO_2_ incubators (Eppendorf, GmBH or Thermo Scientific, USA).

To induce DNA damage, 7 × 10^5^ cell were seeded on 35mm dishes 16 hrs prior to treatment. MG132 and cycloheximide treatments were for 4hrs, Chloroquine for 8 hrs, HDACi, ATRi and UV treatments were for 12 hrs, camptothecin for 16 hrs and etoposide for 24 hrs with concentrations mentioned except for time dependent experiments.

### Plasmid Transfection

RING-finger E3 ubiquitin ligase screen was performed in MCF7 cells as described previously (25). For transfections, 6 × 10^5^ cells were seeded in 35mm dishes and incubated at 37°C overnight. 2 μg of Api5 K251 mutants or Api5 S138A plasmids which were generated by site directed mutagenesis were transfected in MCF7 cells using Lipofectamine 2000 (Thermo Fisher Scientific, USA) using manufactures protocol.

### Generation of stable cell lines

MCF7 cells stably expressing mCherry-tagged Api5 were generated using lentiviral mediated transduction as mentioned previously (bioRxiv; doi: https://doi.org/10.1101/2020.11.22.393256). MCF7 cells stably expressing pLKO1.eGFP and shApi5.eGFP was generated using lentiviral transduction. shApi5 was cloned into pLKO1.eGFP vector as per Addgene protocol using the following primers: Forward Primer: 5’-CCGGAAGACCTAGAACAGACCTTCACTCGAGTGAAGGTCTGTTCTAGGTCTTTTTTTG-3’, Reverse Primer: 5’-AATTCAAAAAAAGACCTAGAACAGACCTTCACTCGAGTGAAGGTCTGTTCTAGG TCTT - 3’. Briefly, 7.5 × 10^5^ HEK293T cells were seeded on 35mm dish and transfected with 1 μg of pLKO1.eGFP or shApi5-eGFP plasmid along with 1 μg of psPAX2 and 0.5 μg pMD2 packaging plasmids using Lipofectamine 2000. 1 ml DMEM containing 30% FBS was added to the cells 24 hours post transfection. 5 × 10^5^ MCF-7 cells were seeded on 35 mm dish for transduction. Viral supernatant was collected 48 hours post transfection and filtered through a 0.45 μm filter to get rid of the cell debris. Filtered viral supernatant containing media along with 1 ml fresh media was added to the MCF-7 cells. 4 μg polybrene was added to the cells to increase the transduction efficiency. Cells were replenished with fresh medium 48 hours post transduction. Transduced MCF7 cells were passaged and sorted using BD Aria Fusion sorter (BD Biosciences, USA). Lentiviral particles containing mCherry-tagged siRNA resistant Api5 wild type and K251 mutants were generated as described previously (bioRxiv; doi: https://doi.org/10.1101/2020.11.22.393256). shApi5 cells after sorting was transduced with lentiviral particles containing mCherry-tagged siRNA resistant Api5 wild type and K251 mutants and sorted for mCherry fluorescence. HeLa cells expressing shFBXW2 were generated as described previously (26).

### MTT assay

For the MTT assay, 1 × 10^4^ cells were seeded in each well of a 96 well plate. 16 hours post seeding, cells were treated with different concentrations of camptothecin and etoposide and incubated for 24 hours at 37°C. After removing the drug containing medium, 0.5 mg/ml thiazolyl blue tetrazolium (MTT) containing DMEM was added to the cells and incubated at 37°C after covering with aluminum foil. 4 hours post MTT addition, MTT-DMEM medium mixture was aspirated from the wells and 100 μl of DMSO was added to dissolve the purple MTT-formazan crystals. The absorbance was recorded at 570nm using the multimode Varioskan Flash plate reader (Thermo Scientific, USA).

### Immunoblotting and Immunoprecipitation

mCherry-tagged Api5 over-expressing stable cells with or without treatments were lysed in cell lysis buffer containing 50 mM Tris-HCl, pH 7.4, 0.1% Triton X-100, 5 mM EDTA, 250 mM NaCl, 50 mM NaF, 0.1 mM Na_3_VO_4_ and protease inhibitors.

Immunoprecipitations were performed as described earlier (27) using Api5, Myc, GFP and ATR specific antibodies. Western blotting was performed with Api5, GFP, ATR, Ubiquitin, pSerine and GAPDH antibodies as previously mentioned (28). Immunoblots include all the experimental lanes of a said experiment. There are no composite images where two or more blots were merged together. The blots were processed as horizontal cut strips in the respective antibody dilutions and imaged as individual strips in ImageQuant LAS4000 (GE Healthcare, now Cytiva, USA).

### Semi-quantitative PCR

RNA extraction and cDNA synthesis of camptothecin, etoposide and UV treated cells were performed as described previously (29). Semi-quantitative PCR was performed using API5 specific primers were described previously (bioRxiv; doi: https://doi.org/10.1101/2020.11.22.393256). GAPDH was used as endogenous control.

### Immunofluorescence

For immunofluorescence, 6 × 10^5^ cells were seeded on coverslips placed in 35mm dishes. After treatment, cells were fixed in 4% paraformaldehyde and permeabilised using 0.5% Triton X after PBS washes. After PBS-Glycine wash, cells were blocked using 10% FBS for 1 hr and incubated overnight with Api5 or pRPA antibody at 4°C. Cells were incubated with secondary antibodies conjugated with Alexa Fluor for 1 hr at room temperature along with phalloidin. Nuclei was stained using Hoechst 33258 for 5 mins and washed twice using PBS. The cells were mounted using mounting media and imaged under 63X oil immersion objective of the SP8 confocal microscope (Leica, Germany).

### Statistical analysis

Densitometry analysis of western blots were performed using ImageJ software. Results were analysed from a minimum of three independent experiments and plotted using GraphPad Prism (GraphPad Software, La Jolla, CA, USA). Student’s t-test and Mann Whitney U test was used to test the significance in difference of Api5 levels between two samples and One-way Anova, Dunnett’s multiple comparisons or Tukey’s multiple comparisons test for more than two samples. p value > 0.05 was considered non-significant. *, ** and *** correspond to p<0.05, p<0.001, and p<0.001 respectively.

## Acknowledgments

We thank Drs Richa Rikhy, and Nagaraj Balasubramanium (IISER Pune, India) for their useful suggestions. The author would like to acknowledge IISER Pune Microscopy Facility and IISER-BD Flow cytometry Facility for access to equipment and infrastructure. We acknowledge assistance from Rintu Umesh and Aishwarya Venkataravi (IISER, Pune) with stable cell line preparation and for designing the schematics in Figure 8 respectively. We also thank Lahiri lab members for their helpful comments and discussions.

## Funding

This study is supported by a grant from Science and Engineering Research Board (SERB), Govt. of India (EMR/2016/001974) and partly by IISER, Pune Core funding. V.K.S was funded by IISER Pune Core funding, S.I was funded by DBT fellowship, J.B by KVPY fellowship, D.P and S.M by DST-INSPIRE Scholarship.

## Conflict of Interest

The authors declare that they have no competing interests.

## Author Contributions

V.K.S, M.K.S and M.L conceived and conceptualised the project. V.K.S performed and analysed the experiments with contribution from S.I, J.B, S.M and D.P. M. K.S gave valuable comments on the first draft of the manuscript. V.K.S and M.L wrote the paper.

## Notes

### Competing Interest Statement

The authors have declared no competing interest.

