## Supplementary Information for "ATR facilitates the degradation of Api5 through the ubiquitin-proteasome pathway via FBXW2 to regulate apoptosis upon DNA damage"

**Supplementary Figure S1**

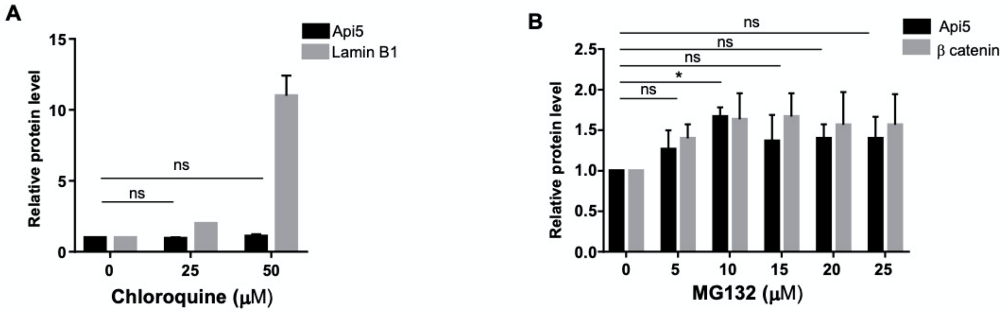


**Figure S1**: Api5 levels of MCF7 cells treated with varying doses of (A) Chloroquine and (B) MG132 for 8hrs were quantified and normalised to GAPDH.

**Supplementary Figure S2**

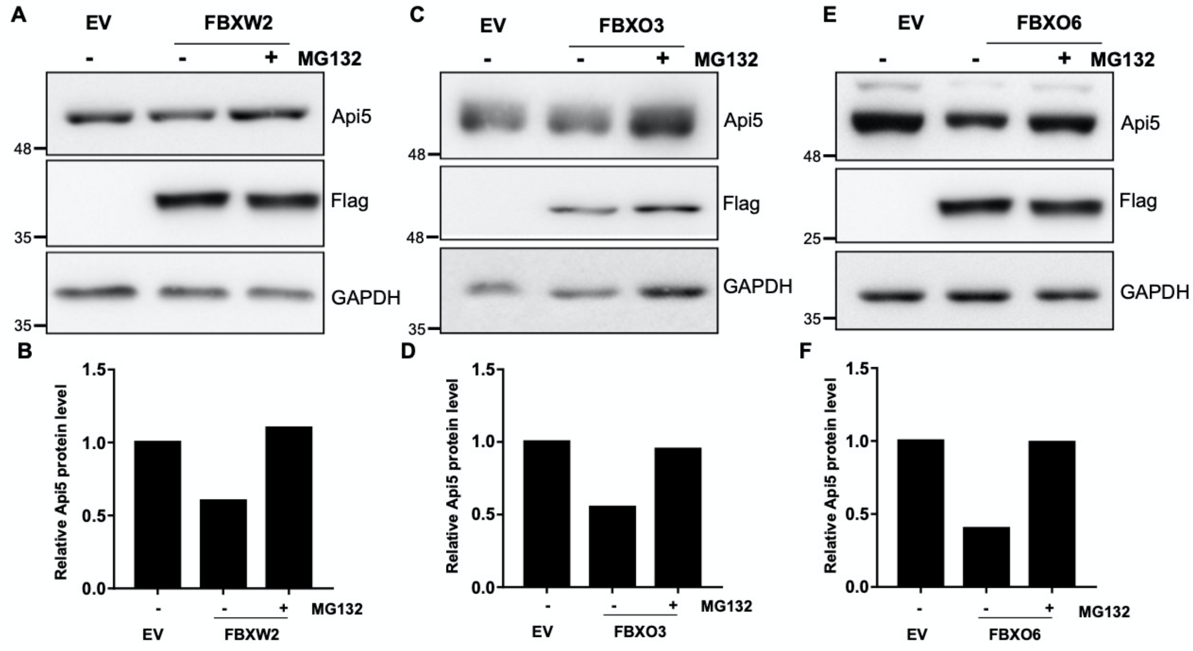


**Figure S2:** MCF7 cells transfected with 1 μg of plasmid containing (A) FBXW2, (C) FBXO3 and (E) FBXO6 were treated with10 μM of MG132 for 4 hrs post 24 hrs of transfection were lysed and Api5 levels were analysed using immunoblotting. (B, D and F) Api5 levels were quantied and normalised to GAPDH.

**Supplementary Figure S3**

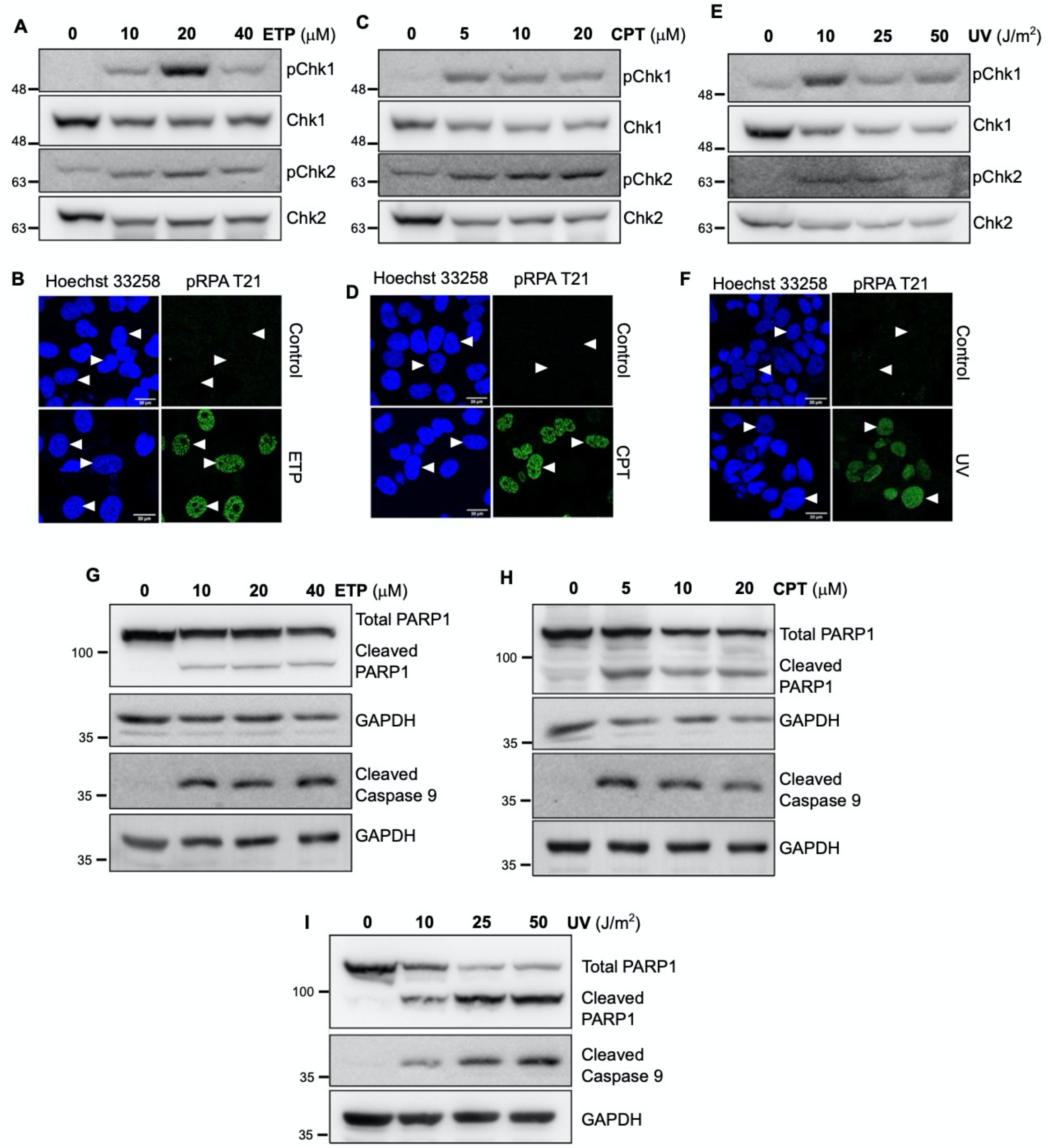


**Figure S3:** MCF7 cells were treated with varying doses of (A and G) etoposide for 24hrs, (C and H) camptothecin for 16 hrs and (E and I) UV for 12hrs were lysed and immunoblotting was performed to analyse the protein levels of pChk1, Chk1, pChk2, Chk2, PARP, and caspase 9. Immunofluorescence was performed to check for RPA activation after (B) 40 μM etoposide for 24hrs, (D) 20 μM camptothecin for 16hrs, and (F) 50J/m$^2$ UV for 12hrs treatment.

### Supplementary Figure S4

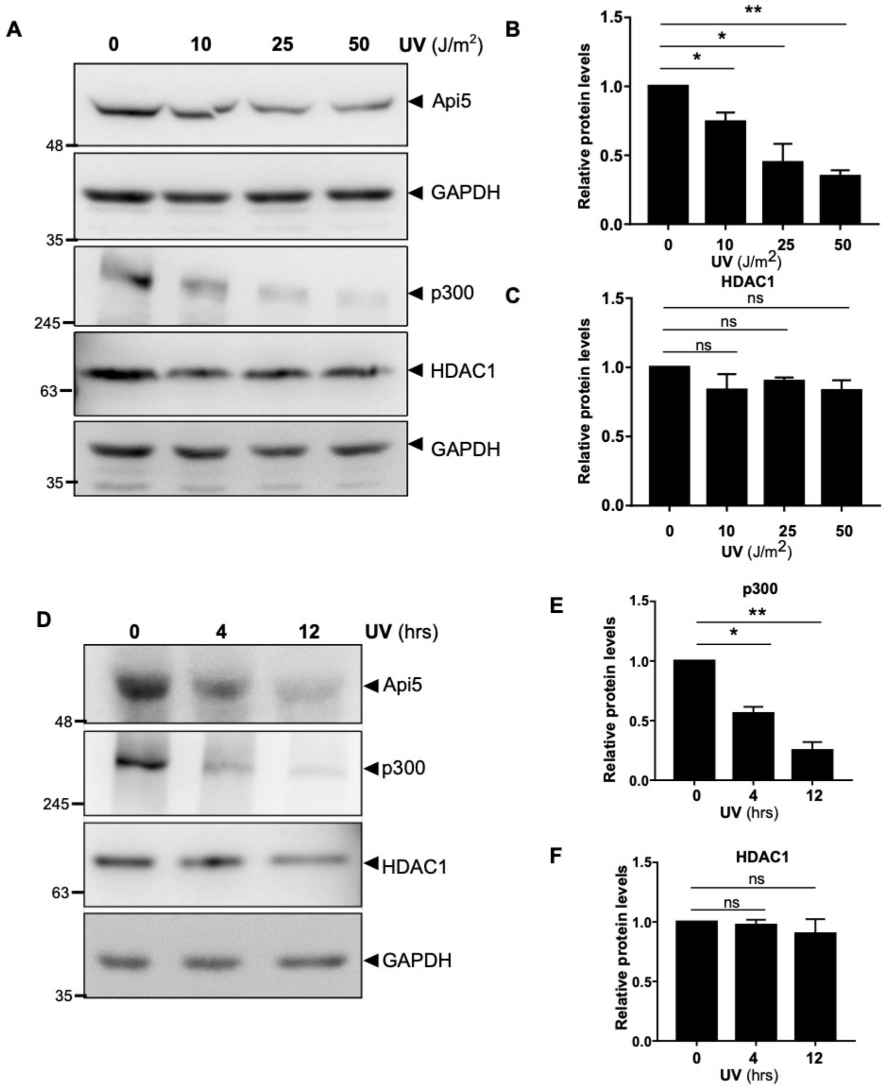


**Figure S4:** MCF7 cells irradiated with (A) varying UV doses for 12 hrs and (D) 50 J/m$^2$ UV for different time points were lysed and immunoblotted to analyse for the levels of Api5, p300 and HDAC1. (B, C, Eand F) protein levels were quantifed after normalising to GAPDH.
